# Number, order and time influence how the brain integrates distinct experiences

**DOI:** 10.64898/2026.09.15.751783

**Authors:** Alina B. Thomas, Francesca S. Wong, A. Simon Killcross, Fred Westbrook, Nathan M. Holmes

**Affiliations:** School of Psychology, University of New South Wales, Sydney, Australia

## Abstract

How does the brain integrate experiences that are separated in time? The present study addressed this question using sensory preconditioning protocols in rats. In these protocols, rats integrate an A-B association (e.g., tone-light) formed in stage 1 with a B- shock association (e.g., light-shock) formed in stage 2 to generate fear responses (freezing) when tested with A alone in stage 3. Here we show that the mechanism of integration depends on the number, order and timing of events across the two stages of training. When the events are novel (low number of exposures) and the interval between stages 1 and 2 is short (24 hours), the A-B and B-shock associations are integrated through formation of a mediated A- shock association during stage 2. By contrast, when the events are more familiar (greater number of exposures), ordered in a particular way, and the interval between stages 1 and 2 is long (14-days), the A-B and B-shock associations are integrated through their chaining at the time of testing with A alone. Thus, number, order and time determine how distinct experiences are integrated in the brain. These findings are discussed with respect to theories of integration and information processing in the medial temporal lobe.

## Introduction

Memory is the means by which the past connects with the present to guide our interactions with the environment. It achieves this feat by integrating the sensory, emotional, spatial and temporal elements of experience. This integration is consistent with a view of memory as a series of episodes that are organized in time ^(1–5)^. However, the memory system can also integrate the elements of experiences that occur days, weeks or months apart. This integration enables one of the most remarkable features of the brain: the ability to infer relations between events that have not (yet) been experienced together. Such inferences allow people and animals to respond appropriately in new situations and solve novel problems. For example, after being attacked by a boy at school, a child may avoid places where the boy had been previously encountered (e.g., the shopping mall); and after learning the relationship between a particular smell (e.g., ripe fruit) and food, a hungry animal may search for food in places where it had previously encountered that smell.

While memory research has largely focused on how discrete episodes are encoded and stored in the brain, less is known about how the brain integrates experiences that are separated in time to permit inferences about unknown relations. In the laboratory, this question can be addressed using sensory preconditioning protocols which exist in a range of species including mice, rats, monkeys and people ^(6–22)^. In a standard sensory preconditioning protocol, subjects are first exposed to pairings of two affectively neutral stimuli (e.g., A→B) in the absence of any other events. Some days later, subjects are exposed to pairings of one of these stimuli (B) with a motivationally significant event (an unconditioned stimulus [US]: appetitive [e.g., food] or aversive [e.g., electric shock]). At test, responding to the other stimulus (A) reveals that animals integrate the A-B association formed at time 1 with the B→US association formed at time 2 to generate responses to a stimulus that was never directly paired with the US.

The integration of experiences in sensory preconditioning can occur through at least two distinct mechanisms ^(9–14,23–27)^. The first is through mediated learning across conditioning of B. The idea is that B activates the memory of its associate A, resulting in formation of an A-US association ^(9–14,23–27)^. Hence, test presentations of A elicit an expectancy of the US and responses in anticipation of its arrival (Figure 1). The second mechanism involves chaining of the trained associations at the time of testing with A: that is, test presentations of A alone generate an expectancy of its associate B which, in turn, generates an expectancy of the US ^(9–14,23–31)^. Hence, A elicits responses via the linking together of the A→B and B→US associations.

**Fig 1.**
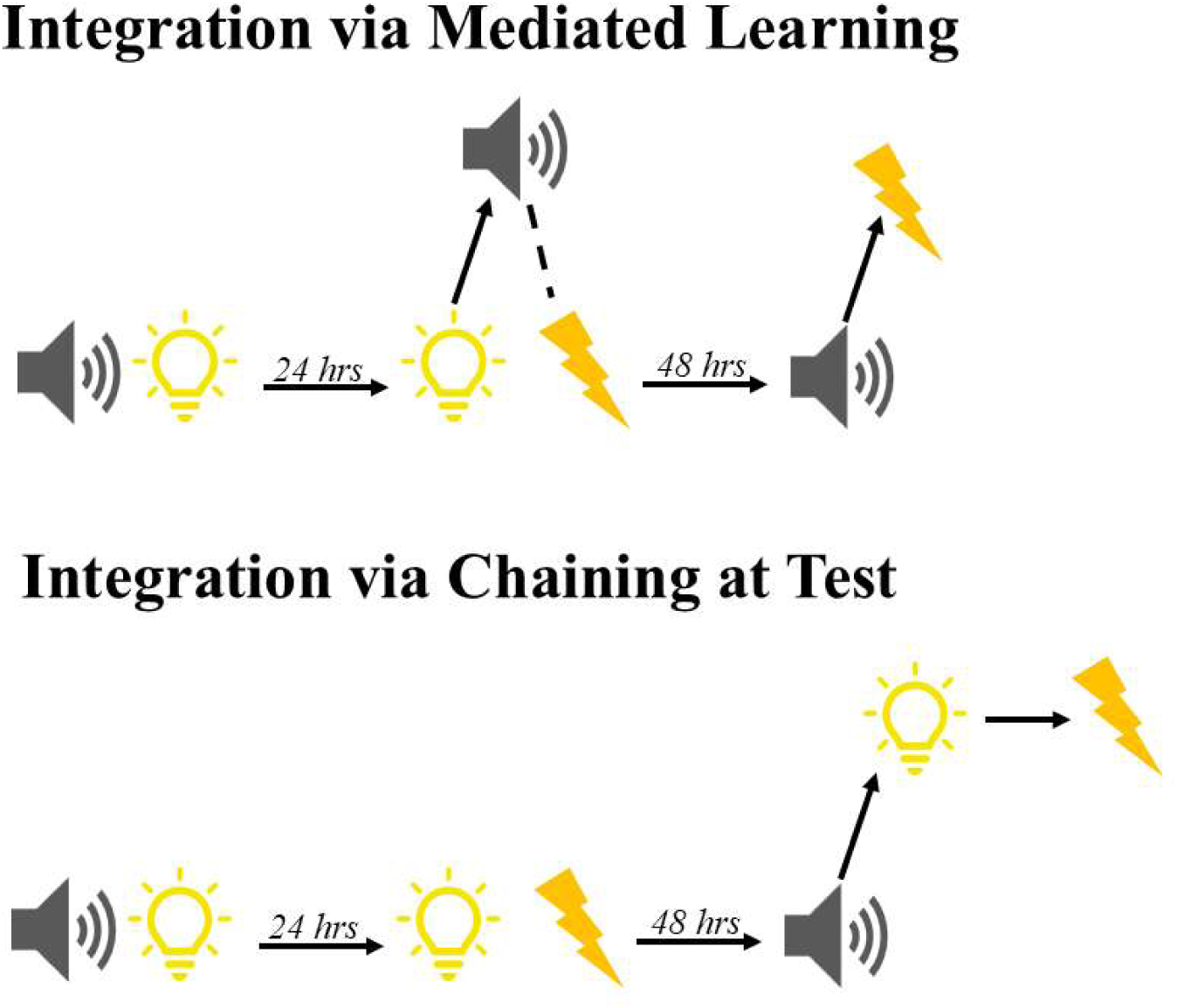
Integration of stage 1 and stage 2 associations in sensory preconditioning. Schematics of how animals could integrate the tone-light (A-B) and light-shock (B-shock) associations to generate fear of the tone (A). **Top panel:** Integration via mediated learning. Here, when animals are exposed to B-shock pairings in stage 2, the presence of stimulus B activates the memory of its past-associate stimulus A, allowing for the formation of a mediated A-shock association. This association is then retrieved at test, resulting in the expression of fear to A. **Bottom panel:** Integration via chaining at test. Here, the A-B and B- shock associations are integrated at the time of testing. When stimulus A is presented at test, animals retrieve the A-B association formed in stage 1 and integrate it with the B-shock association formed in stage 2, resulting in the expression of fear to A.

The fact that integration can occur in either of two ways raises the question: what determines whether integration occurs via one mechanism or the other? The answer to this question is important as the two mechanisms of integration have different implications for how information is represented in memory. Mediated learning implies restructuring of information in memory to account for learning about events that are not physically present; that is, the recoding of A from stage 1 of sensory preconditioning to accommodate the A→US association that forms in stage 2 ^(10,11,23)^. By contrast, chaining at test implies the retrieval and use of past experiences (A→B and B→US pairings) without informational restructuring as it does not involve new learning ^(10,11,23)^. Hence, understanding the factors that promote one form of integration or the other is critical for understanding how information is structured in memory and, importantly, when it is restructured to reflect events that are likely but yet-to-be experienced.

Here, we propose that the mechanism of integration is determined by fundamental properties of experience, such as number, order and time. Specifically, we use the sensory preconditioning protocol described above to test the hypotheses that:

1. mediated learning is more likely when the experiences to be integrated are relatively novel (low number of exposures) ^(27, 35)^.
2. chaining is more likely when the experiences to be integrated are relatively familiar (greater number of exposures) and ordered in a manner that promotes encoding of the associative links that are needed for this form of integration, e.g., A followed by B (A→B) instead of B followed by A (B→A) in stage 1^(26–36)^; and,
3. the mechanism by which two experiences are integrated also depends on the amount of time that has elapsed between them, as well as reminders of experience 1 immediately prior to experience 2.

We test these hypotheses by exploiting what is known about the neural substrates of the associations that form across training in the sensory preconditioning protocol.

Specifically, within the medial temporal lobe, the perirhinal cortex (PRh) receives and processes different types of sensory information, encodes the A→B association that forms in stage 1 of sensory preconditioning but does not encode the B→US association that forms in stage 2 ^(10,11,37)^. Accordingly, we investigate how number, order and time influence the mechanism of integration by blocking activation of *N-methyl-D-aspartate* (NMDA) receptors in the PRh immediately prior to stage 2 of protocols in which we vary the number of A→B or B→A pairings in stage 1, and the interval between stages 1 and 2 ^(38–44)^. If integration occurs through formation of a mediated A→US association at the time of the B→US pairings in stage 2, the PRh manipulation in this stage should disrupt responding to A at test by interfering with retrieval of the information about A and/or its association with the US. By contrast, if integration occurs through chaining of the A→B and B→US associations when A is tested, the PRh manipulation in stage 2 should have no effect on responding to A as rats enter testing with both to-be-chained associations intact.

## Results

In all experiments, the levels of fear responses (freezing) in the experimental chambers (context) prior to stimulus presentations in conditioning and test sessions were low (< 10%) and did not significantly differ between the groups. All diagrams show the mean (±SEM) levels of freezing during acquisition of fear to the light (stimulus B) and test presentations of the tone alone (stimulus A). There was no difference in the acquisition of fear to stimulus B in any experiment.

### Does the mechanism of integration in the “A→B” sensory preconditioning protocol change as a function of the number of stimulus pairings in stage 1?

We have previously shown that, when rats are exposed to just eight A→B pairings in stage 1, the integration of A→B and B→US associations is achieved through formation of a mediated A→US association in stage 2. Here, we hypothesized that increasing the number of A→B pairings in stage 1 would shift the mechanism of integration from mediated learning in stage 2 to chaining at test in stage 3. This hypothesis was based on the idea that increasing the number of A→B pairings should *minimize* reactivation of A in stage 2 (as B is consistently followed by nothing in stage 1) while preserving the associative links needed for chaining at test.

Rats underwent surgery for bilateral cannula implantation in the PRh. After recovery, they were trained in the A→B protocol with either eight or 32 stimulus pairings in stage 1. Immediately prior to the session of B-shock pairings in stage 2, rats received a PRh infusion of the NMDA receptor antagonist, DAPV or its vehicle only (artificial cerebral spinal fluid [ACSF]). Finally, all rats were tested for their levels of freezing during test presentations of A alone. If integration occurs through formation of a mediated A-shock association in stage 2, the DAPV infusion should disrupt freezing to A at test by impairing acquisition of the requisite information in stage 2. If, however, integration occurs through chaining of the A→B and B-shock associations during testing with A alone, the DAPV infusion in stage 2 should have no effect on freezing to A as it spares acquisition of the to-be-chained associations^[10,11]^.

#### Results

The stage 2 DAPV infusion into the PRh reduced freezing to test presentations of A when rats received eight A→ B pairings (Fig 2: *F _1,33_=* 34.8; *p* = 1.29e^-6^; 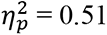, 95% CI: [0.610, 1.253]). This same effect was not seen when rats received 32 A→ B pairings (*F _1,33_=*1.479, *p*= 0.233). This suggest that the mechanism of integration in the A→B protocol changes as a function of the number of stimulus pairings in stage 1. When rats are exposed to just a few A→B pairings in stage 1, integration of the A→B and B→US associations occurs in stage 2 and takes the form of a mediated A→US association ^(10,11)^; hence, among rats exposed to eight A→B pairings in stage 1, blocking NMDA receptors in the PRh prior to the session of B-shock US pairings in stage 2 disrupted freezing to A at test. By contrast, when rats are exposed to many A→B pairings in stage 1, the integration of A→B and B→reinforcer associations occurs via their chaining at the time of testing: hence, among rats exposed to 32 A→B pairings in stage 1, blocking NMDA receptors in the PRh prior to the session of B- shock pairings in stage 2 had no effect on freezing to A at test.

**Figure 2.**
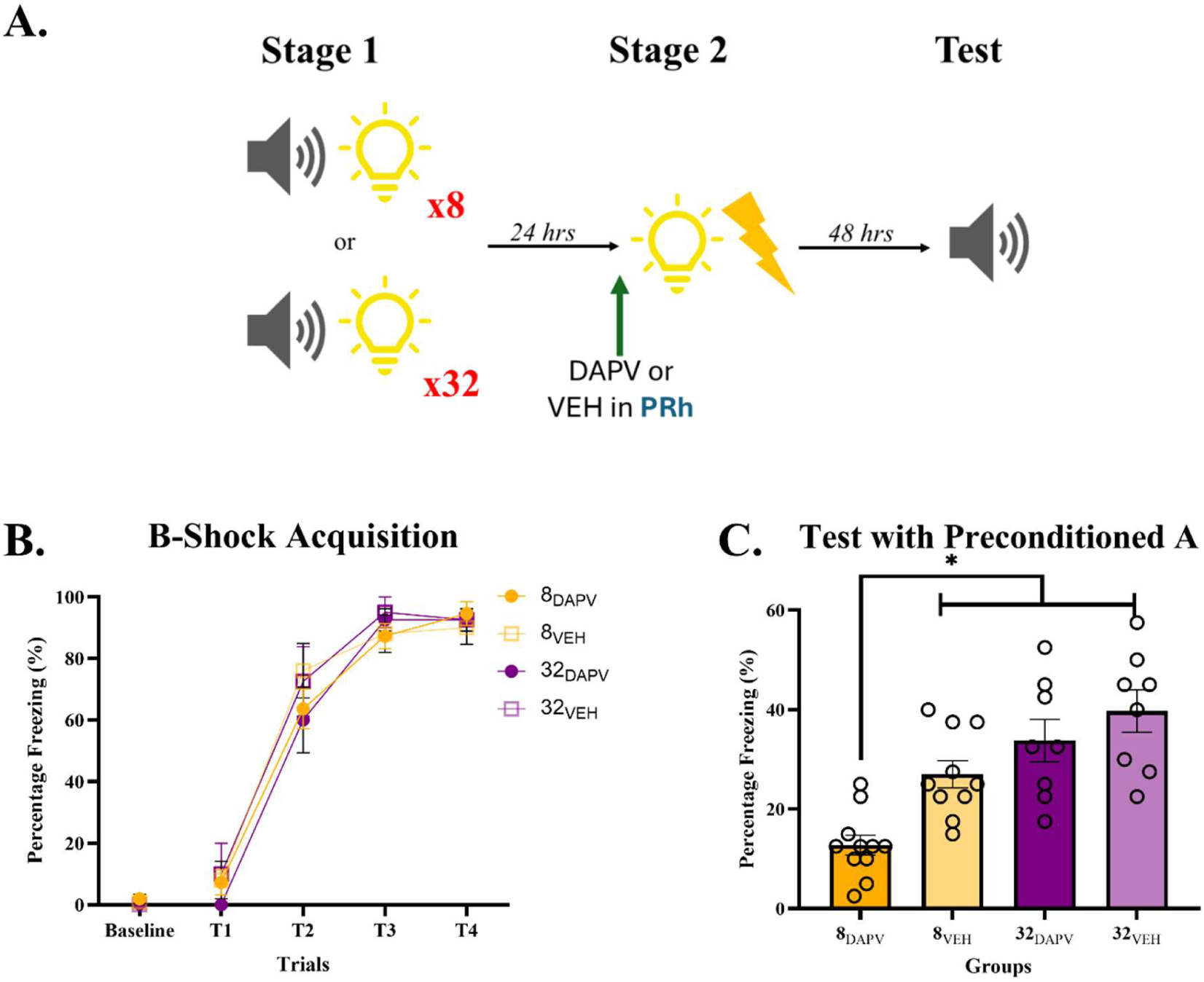
Increasing the number of pairings in stage 1 changes the mechanism of integration in the “A→B” sensory preconditioning protocol. A) Schematic of the behavioral protocol involving *few* A-B pairings in stage 1 and a DAPV (*n*=11) or VEH (*n*=10) infusion prior to stage 2, or involving *many* A-B pairings in stage 1 and a DAPV (*n*=8) or VEH (*n*=8) infusion prior to stage 2. B) Acquisition of freezing across B→shock pairings in stage 2 (*p* = 8.933e^-17^ for linear trend). C) The mean (± SEM) levels of freezing to the preconditioned A across test trials after few or many A-B pairings in stage 1. The symbol ‘*’ indicates a statistically significant difference between the indicated groups (*p* = 1.29e^-6^).

### Why does increasing the number of A→B pairings in stage 1 shift the mechanism of integration from mediated learning in stage 2 to chaining at test in stage 3?

One explanation for this shift is that, as the number of A→B pairings increases in stage 1, rats gradually learn the order in which the stimuli are presented: *“A is followed by B; and B is NOT followed by A.”* Hence, presentations of B no longer reactivate the memory of A during conditioning, resulting in a loss of mediated learning. Instead, presentations of A strongly reactivate the memory of B at test, which serves to complete the chain A→B→shock. Accordingly, we examined whether the shift from mediated learning to chaining at test is conditional on acquisition of this type of order information in stage 1. We did so by examining the likelihood of a shift in the “B→A” sensory preconditioning protocol where rats acquire the opposite order information in stage 1: *“B is followed by A; and A is NOT followed by B.”* If our order hypothesis is correct, an increase in the number of B→A pairings in stage 1 should preserve the likelihood of mediated learning by *strengthening* reactivation of A in stage 2 and undermining the A→B link needed for chaining at test.

Hence, blocking NMDA receptors in the PRh prior to the session of B-shock pairings in stage 2 should disrupt freezing to A at test irrespective of the number of B→A pairings in stage 1.

Rats underwent surgery for bilateral cannula implantation in the PRh. After recovery, they were trained in the B→A sensory preconditioning protocol with either eight or 32 stimulus pairings in stage 1. Immediately prior to the session of B-shock pairings in stage 2, rats received a PRh infusion of the NMDA receptor antagonist, DAPV or its vehicle only (ACSF). Finally, all rats were tested for their levels of freezing during test presentations of A alone.

#### Results

The stage 2 DAPV infusion into the PRh reduced freezing to A at test regardless of whether animals received eight or 32 B→A pairings in stage 1 (Fig 3: *F _1,40_*= 25.788; *p* = 0.00; 5^2^ = 0.40; 95% CI: 0.575, 1.336]). These results show that, in contrast to when the stage 1 pairings are ordered A→B, the mechanism of integration when the order is B→A does *<u>not</u>* change with the number of pairings in stage 1. That is, regardless of the number of B→A pairings in stage 1, integration of the B→A and B→US associations takes the form of a mediated A→US association in stage 2^[10,11]^). Hence, among rats that had been exposed to either eight or 32 B→A pairings in stage 1, blocking NMDA receptors in the PRh prior to the session of B-US pairings in stage 2 reduced freezing to A at test.

**Figure 3.**
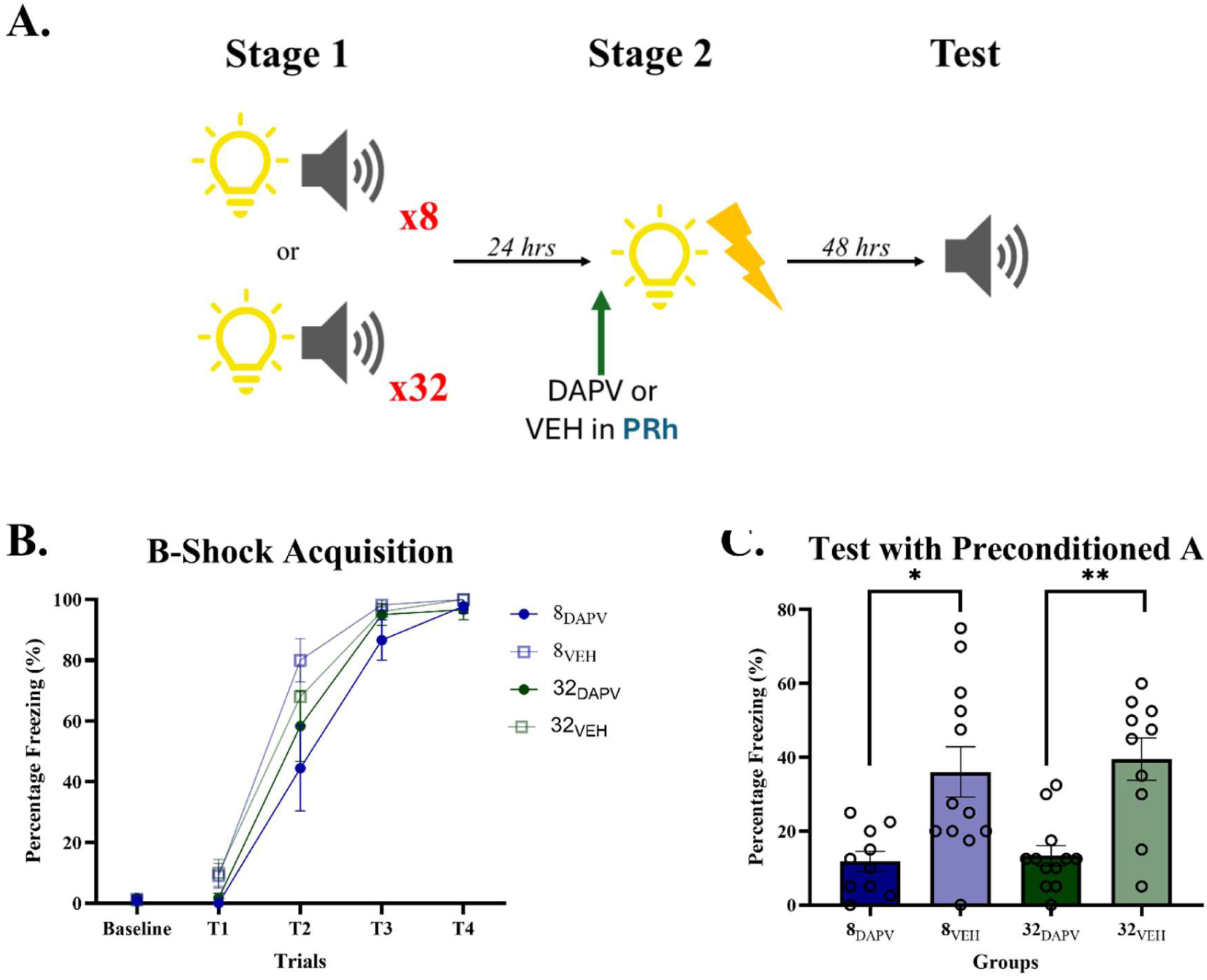
Increasing the number of pairings in stage 1 does not change the mechanism of integration in the “B→A” sensory preconditioning protocol. A) Schematic of the behavioral protocol involving *few* B-A pairings in stage 1 and a DAPV (*n*=12) or VEH (*n*=10) infusion prior to stage 2, or *many* B-A pairings are presented in stage 1 and a DAPV (*n*=12) and VEH (*n*=10) infusion across stage 2. B) Acquisition of freezing across B→shock pairings in stage 2 (*p* < 0.05 for linear trend). C) The mean (± SEM) levels of freezing to the preconditioned A averaged across test trials after few or many B-A pairings in stage 1 (*p* < 0.05). The symbol ‘*’ and ‘**’ indicates a statistically significant difference between the indicated groups (*p <0.05*).

The fact that integration changes in the A→B protocol but not in the B→A protocol shows that it is not simply due to repeated pairings of the A and B stimuli in stage 1: if it was, the same change would have been evident in the B→A protocol. Instead, this shift appears to be due to the rats exposed to many A→B pairings in stage 1 learning that *“A is followed by B; and B is NOT followed by A”*, effectively promoting chaining of the A→B and B→reinforcer associations at test. By contrast, rats exposed to many B→A pairings in stage 1, learn that *“B is followed by A; and A is NOT followed by B”*, which preserves mediated learning in stage 2.

### Is the order information acquired in stage 1 of the A→B sensory preconditioning protocol forgotten with time?

The previous experiments show that when rats are exposed to many A→B pairings in stage 1, the mechanism of integration shifts from mediated learning to chaining at test, and that this shift is due to acquisition of A→B order information in stage 1. The present experiment examined whether the insertion of a 14-day interval between stages 1 and 2 could restore mediated learning. We hypothesized that the 14-day retention interval would cause forgetting of order information acquired across the many A→B pairings in stage 1: hence, presentations of B in stage 2 would reactivate the memory of A, enabling formation of the mediated A-shock association. If this hypothesis is correct, a DAPV infusion into the PRh should have no effect on the test levels of freezing to A when the interval between the many A→B pairings and B-US pairings is just one day, consistent with chaining at test. In contrast, the same infusion should reduce the test levels of freezing to A when the interval between the formation of the two association is 14-days, consistent with mediated learning.

Rats underwent surgery for bilateral cannula implantation in the PRh. After recovery, they were trained in the A→B protocol with 32 A-B pairings in stage 1 and four B-shock pairings in stage 2. For half the rats, the interval between stages 1 and 2 was one day; for the remaining rats, this interval was 14-days. Immediately prior to the session of B-shock pairings in stage 2, rats received a PRh infusion of DAPV or ACSF. Finally, all rats were tested for their levels of freezing during test presentations of A alone.

#### Results

The effect of the stage 2 DAPV infusion on freezing to A at test depended on the interval between stages 1 and 2. Among rats for which this interval was just one day, there was no significant between group differences in freezing to A (*F_1,48_* < 4.043, *p* > 0.05) which replicates the results of the previous experiment. Rats in Group 14D_VEH_ displayed just as much freezing to A as rats in each of the 1-day groups (*F_1,48_* < 4.043, *p* > 0.05), showing that insertion of the 14-day retention interval did not alter the overall level of sensory preconditioned fear to A. By contrast, rats in Group 14D_DAPV_ froze significantly less to A than rats in each of the other groups (Fig 4 **-** *F _1,48_* = 14.801; *p* = 0.000; 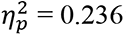; 95% CI: [0.352, 1.123]), showing that the effect of the stage 2 DAPV infusion is, indeed, regulated by the interval between stages 1 and 2.

**Figure 4.**
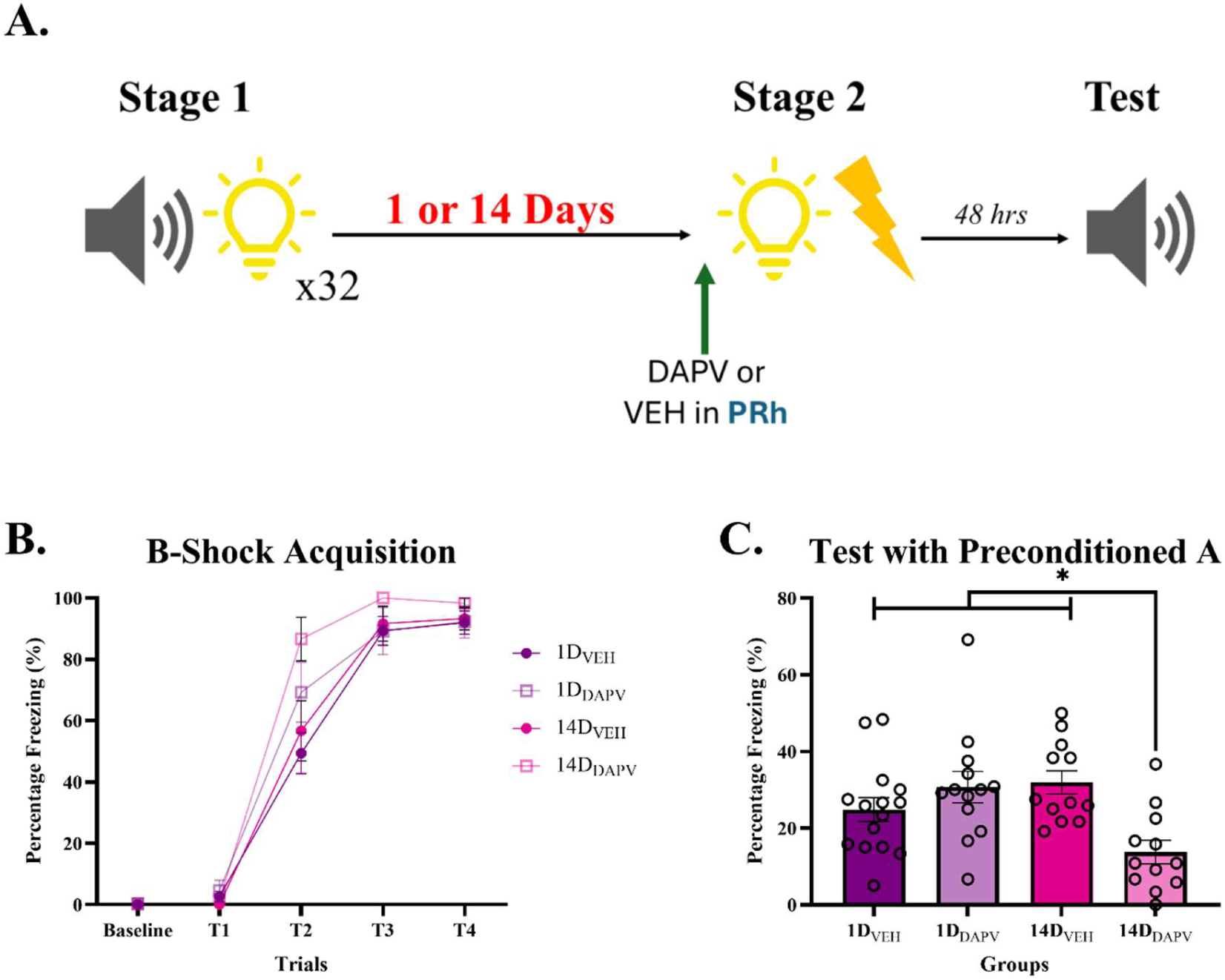
Order information acquired in stage 1 of the A→B sensory preconditioning protocol is forgotten with time. **A)** Schematic of the behavioral protocol used in this experiment. **B)** Acquisition of freezing across B→shock pairings in stage 2 (*p* = 5.421e^-20^ for linear trend). **C)** The mean (± SEM) levels of freezing to the preconditioned A averaged across test trials (*p* < 0.05). The symbol ‘*’ indicates a statistically significant difference between the indicated groups (*p <0.05*).

**Figure 6.**
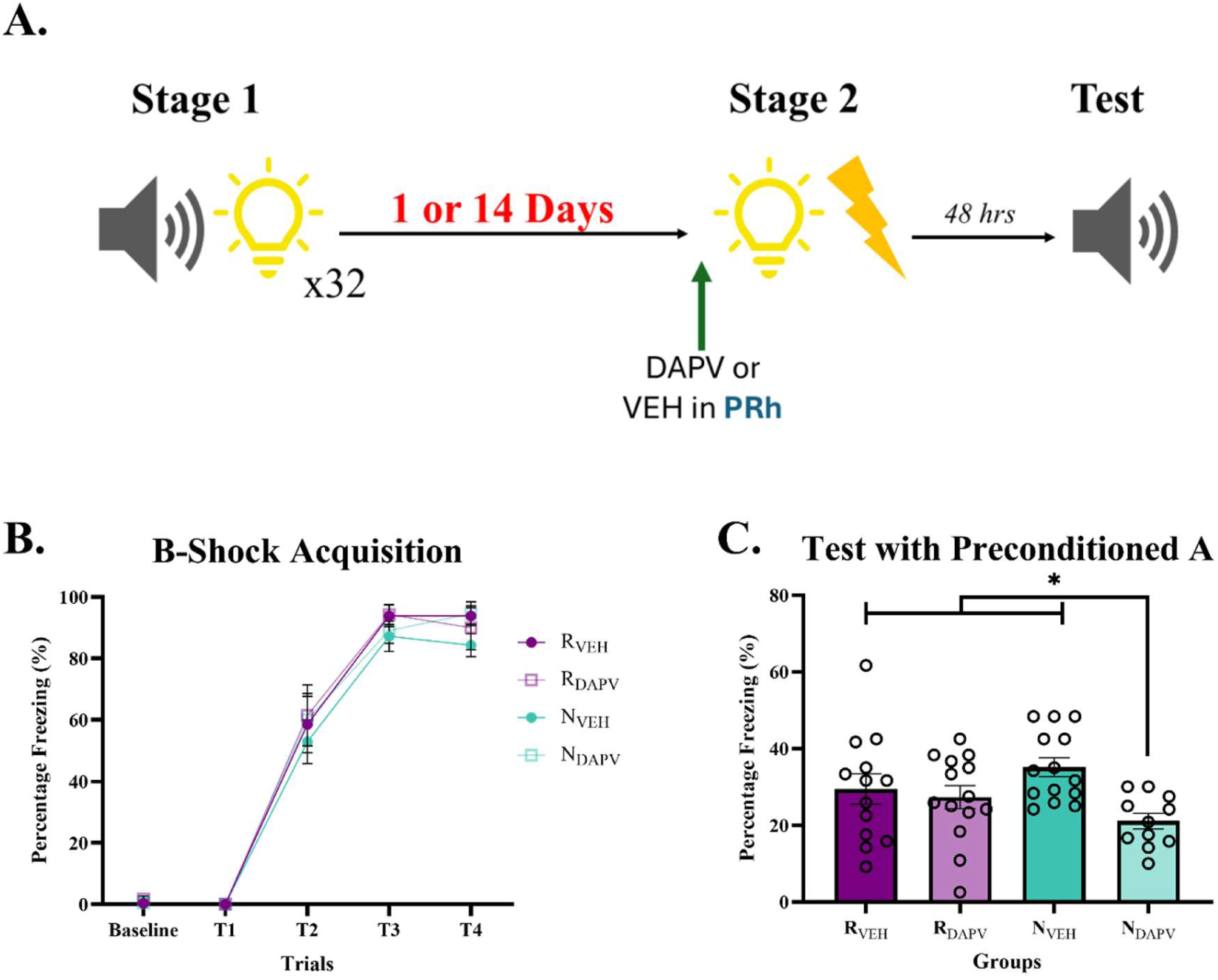
A reminder of the stage 1 experience reverses the effect of time. **A)** Schematic of the behavioral protocol used in this experiment. **B)** Acquisition of freezing across B→shock pairings in stage 2 (*p* = 1.0842e^-19^ for linear trend). **C)** The mean (± SEM) levels of freezing to the preconditioned A averaged across test trials. The symbol ‘*’ indicates a statistically significant difference between the indicated groups (*p* = 0.013).

These results show that, when rats are exposed to many A→B pairings, the mechanism of integration depends on the interval between those pairings in stage 1 and conditioning of B in stage 2. When this interval was just one day, freezing to A at test was unaffected by the DAPV infusion across conditioning, indicating that integration occurred via chaining at test. By contrast, when the interval between the A→B pairings in stage 1 and conditioning of B in stage 2 was 14 days, freezing to A at test was disrupted by the DAPV infusion, indicating that integration was due the formation of a mediated A→shock association.

More generally, these results are consistent with the idea that time preserves the A-B association (a gist-like representation) but prevents retrieval of its specific details ^(45–47)^. The details that fail to be retrieved include the A→B order information (*“A is followed by B; and B is NOT followed by A”*), which naturally shifts the mechanism of integration from mediated learning in stage 2 to chaining at test in stage 3. Hence, the memory of A is once again activated across the B→US pairings in stage 2, resulting in formation of a mediated A→US association that is retrieved and expressed in freezing responses at test.

### Can a reminder of the stage 1 experience reverse the effect of time?

The previous experiment shows that, when rats are exposed to many A→B pairings in stage 1, the shift in the mechanism of integration from mediated learning in stage 2 to chaining at test does not occur when the interval between stages 1 and 2 is increased from 24 hours to 14 days. In other words, the impact of increasing the number of A→B pairings in stage 1 is reversed by the passage of time; and with sufficient time, integration of the A→B and B-shock associations again occurs through formation of a mediated A→shock association in stage 2.

Here, we examine whether this effect of time can be reversed by a reminder of the stage 1 experience immediately prior to stage 2. We hypothesized that an additional session of A→B pairings at the end of a 14-day retention interval would remind rats of their prior stage 1 experiences and, thereby, restore the effects of those experiences on the mechanisms of integration ^(36)^. Specifically, we hypothesized that the A→B reminder session would restore the expectation that *“A is followed by B; and B is NOT followed by A”*: hence, integration of the A→B and B-shock associations would shift back to chaining at test. If this hypothesis is correct, blocking NMDA receptors in the PRh prior to B-shock pairings in stage 2 will have different effects on freezing to A at test depending on the presence versus absence of the stage 1 reminder. Among rats that receive the A→B reminder session prior to stage 2, the DAPV infusion should have no effect on freezing to A at test, indicating successful chaining of the A→B and B-shock associations. By contrast, among rats that receive a session of B→A pairings prior to stage 2, the same infusion should disrupt freezing to A at test, indicating the formation of a mediated A→shock association in stage 2.

Rats underwent surgery for bilateral cannula implantation in the PRh. After recovery, they were trained in the A→B protocol with 32 pairings in stage 1. Thirteen days later, half the rats received an additional, reminder session of eight A→B pairings (reminder [R]) while the remaining rats received a session in which the order of the pairings was reversed, B→A (no reminder [N]). Thus, both groups had the same level of experience with the A and B stimuli but only the former group was reminded of their past stage 1 experience. This was followed by the usual session of B-shock pairings in stage 2 under a PRh infusion of DAPV or ACSF. Finally, all rats were tested for their levels of freezing during test presentations of A alone.

#### Results

The effect of the DAPV infusion on freezing to A at test depended on whether rats were reminded of their stage 1 experience immediately prior to stage 2. The DAPV infusion reduced freezing to A among rats that were *<u>not</u>*reminded of their prior stage 1 experience: rats in Group N_DAPV_ displayed less freezing to A at test than rats in each of the other groups (Fig 5: *F_1,48_* = 6.714, *p* = 0.013, 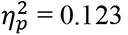; 95% CI: [0.124, 0.984]). Importantly, the reminder had no effect on the overall levels of sensory preconditioned fear as freezing to A in Group N-VEH was the same as freezing to A in Groups R_VEH_ and R_DAPV_ (*F_1,48_* =1.967, p = 0.167). Critically, the DAPV infusion had no effect on freezing to A among rats that *<u>were</u>* reminded of their stage 1 experience: rats in Group R_DAPV_ displayed just as much freezing to A at test as rats in Group R_VEH_ (*F_1,48_* = 0.268, *p* = 0.595).

These results are consistent with the idea that reminding rats of the A→B order facilitates the shift from mediated learning in stage 2 to chaining at test in stage 3. That is, when rats are reminded that “*A is followed by B; and B is NOT followed by A*,” the memory of A is no longer activated across the B-shock pairings in stage 2, which prevents formation of the mediated A-shock association and promotes chaining of the A→B and B-shock associations.

## Discussion

The present study used variations of a sensory preconditioning protocol to identify how rats integrate experiences that are separated in time. It specifically examined how fundamental characteristics of experience such as number, order and time influence the two mechanisms of integration: mediated learning and associative chaining. In each experiment, rats exposed to A-B pairings in stage 1 and B-shock pairings in stage 2 displayed fear responses (freezing) when tested with presentations of A alone. The experiments examined how the mechanism of integration is influenced by changes in the number and order of stimulus presentations in stage 1, the interval between stages 1 and 2, and events that occur at the end of the interval prior to stage 2. We probed the mechanism of integration by blocking activation of NMDA receptors in the PRh immediately prior to the session of B-shock pairings in stage 2. This region encodes the A-B association in stage 1 of sensory preconditioning but plays no role in acquisition of the B-shock association in stage 2 ^(10,11,18)^. Hence, if integration occurs through formation of a mediated A-shock association in stage 2, blocking NMDA receptors in this stage will have disrupted its acquisition and, thereby, freezing responses to A at test. If, however, integration occurs through chaining at test, blocking NMDA receptors in stage 2 will have had no effect on freezing to A at test as each of the to-be-chained associations remains intact ^(10,11)^.

The first major finding in this study was that the mechanism of integration is influenced by the number and order of stimulus presentations in stage 1. When rats were exposed to just a few A→B pairings in stage 1, the stage 2 DAPV infusion into the PRh disrupted the level of freezing to A at test, indicating that integration had been achieved through formation of a mediated A-shock association in stage 2. By contrast, when rats were exposed to many A→B pairings in stage 1, the stage 2 infusion of DAPV into the PRh had no effect on the level of freezing to A at test, indicating that the A→B and B-shock associations were successfully chained upon test presentations of the A alone. Importantly, this shift in the mechanism of integration with a greater number of stimulus pairings was not evident when the order of the stimulus presentations was reversed in stage 1. Amongst rats exposed to either few or many B→A pairings in stage 1, the DAPV infusion in stage 2 disrupted freezing to A at test, indicating that integration typically occurs through mediated learning. Together, these results imply that order information is gradually acquired when the number of stimulus pairings is increased in stage 1 and ultimately determines the mechanism by which the stage 1 and 2 associations are integrated in sensory preconditioning. In the A→B protocol, acquisition of the order information disrupts the likelihood of mediated learning in stage 2 (as B ceases to activate the memory of A) while increasing the likelihood of associative chaining at test (as A strongly activates the memory of B). By contrast, in the B→A protocol, acquisition of the order information preserves the likelihood of mediated learning in stage 2 (as B strongly activates the memory of A) and disrupts the likelihood of chaining at test in stage 3 (as A ceases to activate the memory of B).

The second major finding in the study is that the mechanism of integration is influenced by the interval of time between A→B pairings in stage 1 and B-shock pairings in stage 2. When rats were exposed to many A→B pairings in stage 1 and the interval to stage 2 was just one day, the stage 2 DAPV infusion into the PRh had no effect on the level of freezing to A at test, indicating that the A→B and B-shock associations were successfully chained upon presentations of the A alone. By contrast, when rats were exposed to many A→B pairings in stage 1 and the interval to stage 2 was increased to 14 days, the DAPV infusion in stage 2 disrupted freezing to A at test, indicating that the lapse of time between stages 1 and 2 reinstated mediated learning as the mechanism of integration. Importantly, however, this effect of time was not evident when rats were reminded of their stage 1 experience immediately prior to stage 2. Amongst rats that received an additional session of A→B pairings prior to B-shock pairings in stage 2, the DAPV infusion in stage 2 had no effect on freezing to A at test, indicating that the stage 1 and 2 associations were again integrated through chaining at test. This pattern of results differed from those obtained among rats that received an additional session of B→A pairings on the day prior to the session of B- shock pairings in stage 2: under these circumstances, the DAPV infusion disrupted freezing to A at test, indicating integration occurred through mediated learning. Together, these results imply that the order information which is gradually acquired when the number of stimulus pairings is increased in stage 1 is forgotten when the interval between stages 1 and 2 is increased to 14 days; but can be reinstated by a reminder of the stage 1 experience immediately prior to stage 2. Hence, the interval between stages 1 and 2 as well as reminders delivered at the end of this interval combine to determine the mechanism by which the stage 1 and 2 associations are integrated in sensory preconditioning ^(48)^. It is worth noting that the interval of time between stages 1 and 2 did not cause forgetting of the A-B association *per se*: if it had, there would have been a complete loss of sensory preconditioned fear to A. Instead, time preserved the general features of the audiovisual A-B association but caused forgetting of the specific order information relating to its individual events: hence the shift in the basis of sensory preconditioned fear from chaining at test to mediated learning in stage 2. These dissociable effects of time are consistent with those reported in a variety of domains, including memory for real life and laboratory events in people ^(43,46,48)^.

As noted above, mediated learning and associative chaining have different implications for the way that information is represented in memory. Consistent with such differences, the two types of integration require communication between the PRh and BLA but differ with respect to the precise timing of this requirement. Briefly, when rats are exposed to just a few A→B pairings in stage 1, formation of a mediated A-shock association requires communication between the PRh and BLA across the session of B-shock pairings in stage 2: this is the moment when the memory of A is activated in the PRh and recoded to associate with foot shock in the BLA. It does not, however, require communication between the PRh and BLA across testing with the A alone in stage 3 ^(11)^. By contrast, when rats are exposed to many A→B pairings in stage 1, chaining of the A→B and B-shock associations does not require communication between the PRh and BLA across the session of B-shock pairings in stage 2 but does require this communication across testing with the A alone in stage 3: this is the moment when the A→B association which is encoded and stored in the PRh is retrieved and linked with the B-shock association which is encoded and stored in the BLA ^(11)^. Future work should examine how the PRh and BLA combine to support successful integration under the different circumstances of training and testing described here.

Finally, the present findings have several implications for psychopathologies characterized by disturbances in memory and dysfunction in the medial temporal lobe. These include post-traumatic stress disorder (PTSD) in which fear can be triggered by stimuli that were not present at the time of an aversive experience ^(49–52)^. They also include schizophrenia and schizoaffective disorders (SSDs), which can be characterized by false or delusional beliefs about relations between environmental events ^(44,53,54)^. Specifically, by identifying the factors that determine whether memories are integrated through mediated learning (a form of false belief) or associative chaining, as well as the neural circuitry that underlies the two types of integration in the amygdala and perirhinal cortex, our findings suggest that the overgeneralization of fear in PTSD and delusional beliefs in SSDs may have a common underlying cause in the failure of amygdala-cortical interactions ^(55,56)^. Future work will test this hypothesis by identifying factors that influence the mechanism of integration in neurotypical humans (e.g., memory for the order of events), examining whether sensitivity to these factors is altered among specific clinical populations (e.g., PTSD, SSD), and comparing the neural correlates of change in sensitivity among neurotypical and clinical populations. This work will continue to advance our understanding of how the brain integrates experiences that are separated in time to solve novel problems; and provide insights to psychopathology characterized by disturbances in memory.

## Materials and Methods

### Subjects

The subjects were male and female Long Evans rats. Rats were housed with littermates by sex in plastic boxes (40 cm wide x 22 cm high x 67 cm long) ^(9,10)^. Females were housed in groups of eight and males in groups of three to four. The home cages were kept in a climate-controlled colony room maintained at 21°C and on a 12-hour light/dark cycle (lights on at 0700). Food and water were provided *ad libitum* for the duration of the experiment. Rats were handled for five days prior to the commencement of experiments to familiarize them with the experimenter.

### Apparatus

Experiments were conducted in four identical chambers (26 cm long x 30 cm wide x 30 cm high). Each chamber had plastic front and back walls and aluminium side walls and ceilings. The floor of the chamber was a grid made of stainless-steel rods, each 2 mm in diameter and spaced 15 mm apart from center-to-center^(10,11)^. Below the chamber was a waste tray containing corn cob bedding material, which was cleaned and replaced with fresh bedding after each session. Each chamber was enclosed in a light- and sound-attenuating wooden cabinet with black walls. A speaker mounted on the back wall of the cabinet was used to play an 800 Hz tone at 70 dB above a background noise level of 45 dB. A 2x3 array of LEDs, also mounted on the back wall, was used to deliver a flashing light at a frequency of 3.5 Hz ^(10,11)^. A constant-current generator connected to the grid floor of each chamber was used to deliver a shock of 0.8 mA for 0.5 s. Cameras mounted inside each light- and sound-attenuating cabinet were connected to a computer in an adjacent room and used to record behavior for offline scoring of freezing responses. The tone, light and shock presentations were controlled via the same computer using MATLAB (MathWorks) software.

### Surgery

All rats were surgically implanted with bilateral cannulas targeting the PRh. Rats received 5% isoflurane in oxygen inside an induction chamber until fully anaesthetized (approximately four min). They were then removed and positioned on a stereotaxic apparatus (David Kopf Instruments) and injected with: 1) a local anaesthetic along the line of the first incision (0.5% bupivacaine, subcutaneous [s.c.] injection, 0.2 ml) and 2) a non-steroidal anti-inflammatory for the purposes of pain relief (50mg/ml carprofen, s.c., 0.3 ml)^(10,11)^. The first incision was then made to expose the skull, and two holes were drilled to allow for insertion of two 26-gauge guide cannulas (Plastics One, Inc.) into the PRh at coordinates 4.3 mm posterior to Bregma, 5.5 mm lateral to the midline and 8.4 mm ventral to the skull surface, angled at approximately 9° ^(10,11)^. Cannulas were secured in place with dental cement and jeweller’s screws. A dummy cannula was inserted into each guide cannula to preserve its patency and only removed during drug infusions. Rats received an intraperitoneal (i.p.) injection of procaine penicillin (approximately 0.5 ml of a 300 mg/kg solution) post-surgery to protect against infection. They were placed on a heating mat in a recovery box until fully recovered from anesthesia and then returned to their home cage. All rats were monitored and weighed daily for seven days before behavioral training commenced ^(10,11)^.

### Drugs

The selective NMDAr antagonist, DAPV was dissolved in artificial cerebrospinal fluid (ACSF; Sigma-Aldrich) to yield a final concentration of 10 µg/µL ^(26)^. The drug solution or vehicle (ACSF) alone was infused into the PRh at a rate of 0.25 µL/min for two min, resulting in a total infusion of 0.5 µL into each hemisphere ^(10,11,57)^.

### Drug Infusions

Prior to the infusion day, rats were familiarized with the infusion room and removal of the dummy cannulas. This was done to minimize any effects of the infusion procedure on conditioning. Prior to light-shock pairings, dummy cannulas were again removed and replaced with two 33-gauge infusion cannulas: one in each guide cannula. Infusion cannulas were connected to 25 µL Hamilton syringes via polyethylene tubing; and the syringes were fixed to a programmable infusion pump (World Precision Instruments). The pump was programmed to infuse the drug or vehicle alone into the PRh at a rate of 0.25 µL/min for two min. Upon completion of the two min infusion period, infusion cannulas remained in place for an additional two min to allow for diffusion of the drug or vehicle away from the cannula tip, making a total infusion time of four min. Infusion cannulas were then removed, and dummy cannulas reinserted. Rats were then returned to their home cage for a few min before being placed in the chambers for their training session.

### Histology

At the completion of behavioral training and testing, rats were euthanized with a lethal dose of sodium pentobarbital. Brains were extracted and cut on a cryostat into 40 µm thick coronal sections. Every second section was mounted on a glass microscope slide and stained using cresyl violet. A light microscope was then used to identify the placement of each cannula tip with respect to the PRh using well-defined boundaries ^(57)^. Rats with at least one misplaced cannula were excluded. The placement of correctly positioned cannulas is shown in Fig S6 for all experiments.

### Behavioral Procedure

#### Context Exposure

Rats were placed in conditioning chambers for two 20-min sessions on days 1 and 2. Each session was separated by a fixed interval of three hours (one in the morning and one in the afternoon). This was done to familiarize rats with the chambers and, thereby, reduce subsequent conditioning of freezing to the context.

#### Sensory Preconditioning (Stage 1)

All rats were placed in the context and exposed to either eight presentations of the tone and flashing light in a single session (day 3), or 32 presentations of the tone and light across four daily sessions (days 3-6). All tone presentations were 30 s in duration while all flashing light presentations were 10 s in duration ^(10,11)^. The first stimulus presentation occurred five min after rats were placed in the chamber. In cases where stimuli were presented in a paired manner, the offset of one stimulus co-occurred with onset of the other stimulus, and the interval between each of the pairings was exactly five min. In cases where the stimuli were presented in an unpaired manner, they were presented in an alternating sequence and the interval between the stimulus presentations was exactly 150 s ^(10,11)^. After the final stimulus presentation rats remained in the chamber for an additional min before being returned to their home cages.

#### Reminders

In some experiments, rats were exposed to 32 stimulus pairings in stage 1 and, 13 days later, an additional session of 8 stimulus pairings. The details of this additional session were exactly as described above for sensory preconditioning: the session was intended to remind rats of their prior stage 1 experience.

#### Direct Fear Conditioning (Stage 2)

Rats were placed in the context and exposed to four presentations of the 10 s light, each of which co-terminated with foot shock (0.8 mA, 0.5 s). This occurred on day 4 for rats exposed to eight stimulus pairings in stage 1, day 7 for rats exposed to 32 stimulus pairings in stage 1 and no delay to stage 2, and day 21 for rats exposed to 32 stimulus pairings in stage 1 and a 14-day interval to stage 2. The first light presentation occurred five min after rats were placed into the chambers, the interval between the light-shock pairings was fixed at five min, and rats remained in the chambers for two min after the final light-shock pairing.

#### Context Extinction

The day after direct fear conditioning of the light, rats received two 20-min sessions of context alone exposure spaced three hours apart. The interval between these sessions was spent in the home cages. The sessions were intended to extinguish any freezing to the context alone, which would otherwise obscure that elicited by the stimuli on the subsequent tests.

#### Testing (Stage 3)

Over the next two days, rats were tested for their levels of freezing to the sensory preconditioned tone and the directly conditioned light. On the first of these test days (day 6 for rats exposed to eight stimulus pairings in stage 1, day 9 for rats exposed to 32 stimulus pairings in stage 1 and no delay to stage 2, and day 23 for rats exposed to 32 stimulus pairings in stage 1 and a 14-day delay to stage 2), rats received another 10 min session of context alone exposure to further reduce any spontaneously recovered context freezing. They were then returned to their home cages. Three hours later, they were again placed in the context for testing. This consisted in eight presentations of the tone alone. Each tone presentation lasted for 30 s, the first tone presentation occurred 120 s after placement in the context and the interval between tone presentations was fixed at 180 s. Rats remained in the context for an additional two min after the final tone presentation. On the second of the test days (day 7 for rats exposed to eight stimulus pairings in stage 1, day 10 for rats exposed to 32 stimulus pairings in stage 1 and no delay to stage 2, and day 24 for rats exposed to 32 stimulus pairings in stage 1 and a 14-day delay to stage 2), rats received 12 presentations of the light alone. Each light presentation lasted for 10 s. All other details were as described for testing with the tone alone^(10,11)^.

### Scoring and Statistics

All conditioning and test sessions were digitally recorded for offline scoring of freezing. This was defined as the absence of all movement other than that required for breathing ^(58)^. Rats were scored as ‘freezing’ or ‘not freezing’ every two seconds during periods of interest: a two- or five-min baseline period prior to the first stimulus presentation, and the stimulus presentations themselves. The percentage of time spent freezing in these periods was calculated by dividing the total number of samples scored as freezing by the total number of observed samples. An observer blind to the rat’s group allocation also scored a randomly selected 50% of the training data and all test data. The correlation between the scores obtained by the experimenter and naïve observer was high (Pearson > 0.9). The data were analysed using a set of planned orthogonal contrasts, with the type 1 error rate controlled at α = 0.05 ^(58)^. Standardized 95% CIs are reported for significant differences and either Cohen’s *d* or partial eta-squared ^(52)^ is reported as a measure of effect size (where 0.14 is a large effect size). The required number of rats per group was based on our prior studies of sensory preconditioning which indicated that eight subjects per group provides sufficient statistical power to detect an effect size greater than 0.06 (partial eta-squared) with the recommended probability of 0.8-0.9.

